# A systematic NGS-based approach for contaminant detection and functional inference

**DOI:** 10.1101/741934

**Authors:** Sung-Joon Park, Satoru Onizuka, Masahide Seki, Yutaka Suzuki, Takanori Iwata, Kenta Nakai

## Abstract

**Background:** Microbial contamination impedes successful biological and biomedical research. Computational approaches utilizing next-generation sequencing (NGS) data offer promising diagnostics to assess the presence of contaminants. However, as host cells are often contaminated by multiple microorganisms, these approaches require careful attention to intra- and interspecies sequence similarities, which have not yet been fully addressed.

**Results:** We present a computational approach that rigorously investigates the genomic origins of sequenced reads, including those mapped to multiple species that have been discarded in previous studies. Through the analysis of large-scale synthetic and public NGS samples, we approximated that 1,000−100,000 microbial reads prevail when one million host reads are sequenced by RNA-seq. The microbe catalog we established included *Cutibacterium* as a prevalent contaminant, suggesting that contamination mostly originates from the laboratory environment. Importantly, by applying a systematic method to infer the functional impact of contamination, we revealed that host-contaminant interactions cause profound changes in the host molecular landscapes, as exemplified by changes in inflammatory and apoptotic pathways during *Mycoplasma* infection.

**Conclusions:** These findings reinforce the concept that precise determination of the origins and functional impacts of contamination is imperative for quality research and illustrate the usefulness of the proposed approach to comprehensively characterize contamination landscapes.

## Background

In contemporary biology, cell resources are routinely manipulated via various techniques under a range of conditions. During the course of such manipulations, eukaryotic cells are potentially exposed to microorganisms that cause prominent morphological and physiological changes in their host cells, and such changes often result in erroneous experimental conclusions [1–3]. In medical and clinical settings, it is imperative to detect infectious agents in donated cells to avoid donor-patient disease transmission [4–6]. Despite a community-wide effort to introduce precautions to prevent contamination, the pervasiveness of unexpected microbial contaminants in publications has recently been reported [7–9]. This diminished quality is due, in part, to intrinsic difficulties in assaying for contamination, e.g., window periods, primer dependency, and drug resistance. As an alternative solution to these problems, next-generation sequencing (NGS) has been shown to be an effective approach [6, 10, 11].

Recently, NGS-based studies have intensively addressed the presence of specific microorganisms (e.g., *Mycoplasma*) [7–9] and the influence of cross-contamination caused by exogenous sources (e.g., laboratory reagents and sequencer carryover) [12–15]. While computational methods employing efficient bioinformatics strategies have greatly contributed to such studies [16–19], fundamental challenges still remain [20, 21]. One difficulty in particular is how to deal with sequenced reads that can be mapped to multiple microbial genomes simultaneously, which leads to detection uncertainty [17, 21, 22]. In fact, biological resources contaminated by multiple microorganisms are not uncommon, and the nature of higher intra- and interspecies sequence similarities in microbial communities is well known; that is, distinct species belonging to the same genus have >97% sequence identity [23]. There are also species in different genera that are difficult to distinguish genomically [21]; for instance, the genome sequence of Enterobacteria phage phiX174, a routinely used spike-in species in Illumina sequencing, shares >95% identity with the sequences of the G4 and Alpha3 Microvirus genera [24].

In this study, to improve the certainty of NGS-based contaminant detection, we developed a computational approach that rigorously investigates the genomic origin of sequenced reads. Unlike existing rapid and quasi-alignment approaches, our method repeatedly performs read mapping coupled with a scoring scheme that weights the reads unmapped to the host genome but mapped to multiple contaminant genomes. This approach allows estimation of the probability of chance occurrence of the detected contaminants. By setting human as a host and bacteria/viruses/fungi as contaminants, we demonstrate the robust performance of the proposed method by analyzing synthetic data. Next, we analyzed over 400 NGS samples to profile the contamination landscape, which yielded a catalog of the microbes prevalent in the molecular experiments. Furthermore, we applied a matrix factorization algorithm using our profiles to infer the functional impacts of contamination, thus providing a novel window into the complexities of host-microbe interactions.

## Results

### Identification and quantification of host-unmapped microbial reads

Our first goal was to extract exogenous reads from the input NGS reads by performing greedy alignments. Similar to the initial screening step in published methods [18, 25, 26], our method thoroughly discards host-related reads (Step I to IV in Fig. 1A). Unlike the sequential subtracting approach used in other published methods [13, 18, 25], our method independently maps the screened reads to individual microbial genomes (Step V in Fig. 1A), which enables us to define the mapping status of each read (Step VI in Fig. 1A), i.e., a read is categorized as either a ‘uniq-species-hit’ (or ‘uniq-genus-hit’), which is uniquely mapped to a specific species (or genus), or as a ‘multi-species-hit’ (or ‘multi-genera-hit’), which is repeatedly mapped to multiple species (or genera).

**Figure 1.**
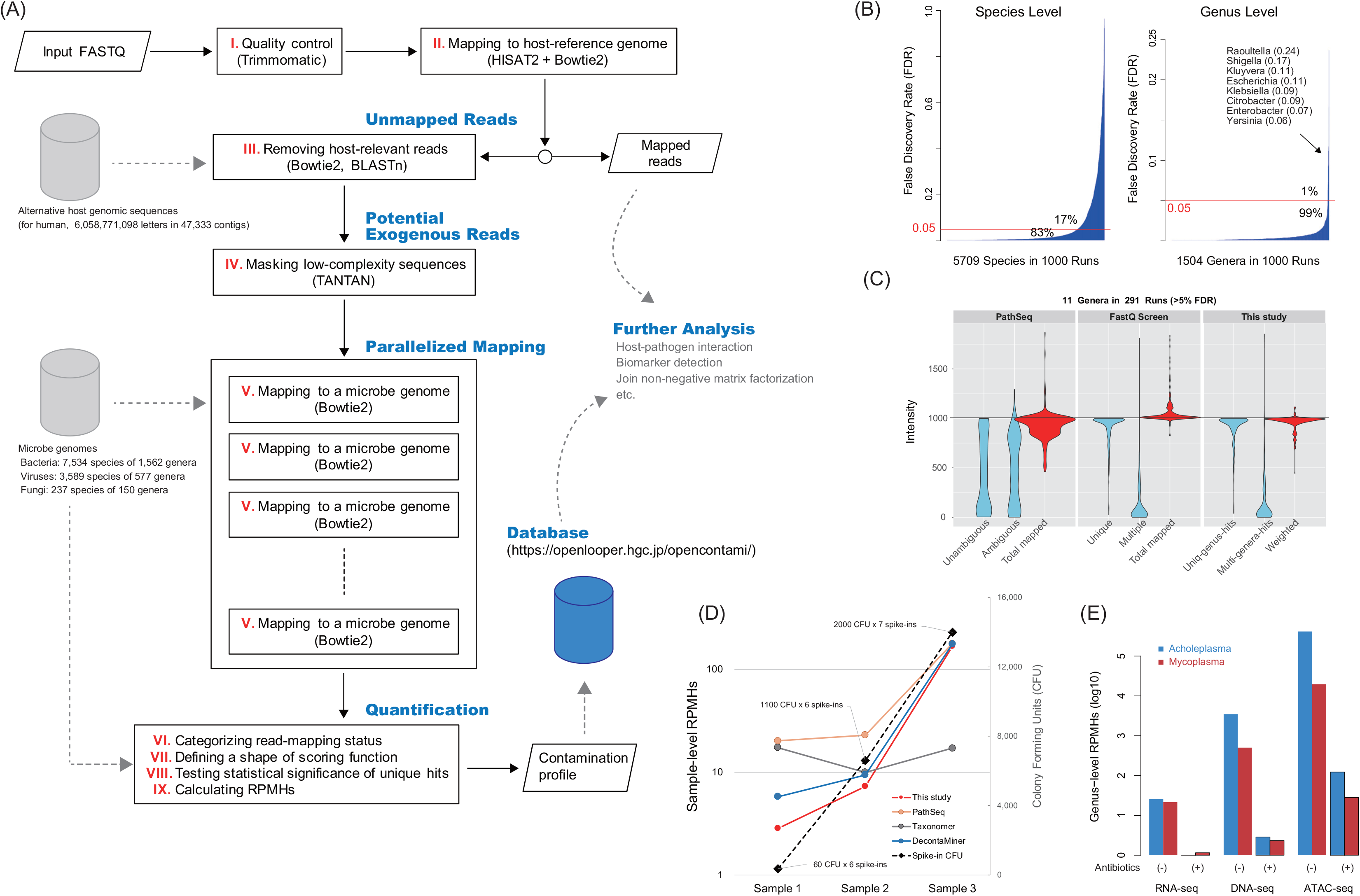
Overall structure of the proposed pipeline and results of the performance assessment. (A) Schematic representation of the proposed pipeline that executes rigorous read alignment with a large-scale genome database. (B) FDR distribution in the reversion tests considering falsely mapped reads to other species or to other genera. Particular genera, including Raoultella, Shigella, and Kluyvera, are difficult to distinguish genomically. (C) Comparative analysis for the effects of uniq-genus-hits and weighted multi-genera-hits in quantification. “Total mapped” represents the sum of uniq-genus-hits (Unique and Unambiguous) and multi-genera-hits (Multiple and Ambiguous). “Weighted” represents the adjusted “Total mapped” by our scoring scheme. (D) Correlations between the detection quantification and spike-in concentration assayed by DNA-seq (0-day cultured hPDL-MSCs with antibiotics). (E) RPMH differences among three NGS protocols in Mycoplasma spike-in detections (3-day cultured hPDL-MSCs).

Prior to quantifying microbe abundance, our method tests the statistical significance of the unique microbe hits by preparing an ensemble of unique hits with random read sets (Step VIII in Fig. 1A). If the observed value of the unique hits is significantly greater than its random ensemble mean value, the pipeline reports the microbe as a potential contaminant. Microbes that were detected with no unique hits are considered not of interest. Next, to calculate an RPMH (read per million host-mapped reads) value for each species (or genus), our method weights the reads repeatedly mapped to the multiple microbes reported (Step VII in Fig. 1A). The RPMH at a sample level is based on the sum of the raw counts of microbe-mapped reads. In summary, the proposed method explores uniquely mapped reads, as a primary key, and exploits the weighted contributions of reads mapped to multiple microbial genomes (see the “Methods” section).

### Parameter tuning with simulated reads

To assess the performance of our mapping approach (Step V and VI in Fig. 1A), we first conducted a reversion test with random microbial read sets, which measures the ratio of reads that correctly mapped to their origin genomes. We prepared 10,000 reads (1,000 ×10 species) per run and repeated the test 1,000 times with different read sets. We also tested different parameters for Bowtie2 [27]. Since the reversion test uses intact DNA fragments randomly selected, if the pipeline works perfectly, all the species will be detected with the 1,000 reads.

With the default parameters (Fig. 1B), when counting false positives at the species level (i.e., multi-species-hits), 17% of the tested species had over 5% multi-species-hits. When allowing reversion errors within the same genus (i.e., counting uniq-genus-hits), only 0.7% of the genera (11 out of 1504) showed over 5% multi-genera-hits. The other parameters of Bowtie2 had no effect on these results (Additional file 1: Fig. S1A-C). This observation implies the presence of high sequence similarity at the species level. We calculated the ratios by running PathSeq [18], FastQ Screen [28], and DecontaMiner [29] (Additional file 2). Of note, comparing existing pipelines is not straightforward because different aligners are employed and databases are inaccessible in some cases. With this in mind, the results indicated that the pipelines exhibit inferior performance for a portion of the reads, similar to our pipeline (Additional file 1: Fig. S2A). These results suggest that the FDRs likely depend on the degree of microbial intra-species sequence homology causing ambiguous multi-species-hits, rather than on intrinsic algorithmic differences in the pipelines.

We next investigated the influence of interspecies sequence homology. Overall, although the reversion test ensures 1,000 microbial reads as the intensity of a species, counting only the uniq-genus-hits showed lower intensity (i.e. loss of accuracy due in part to the occurrence of multi-genera-hits), while taking the sum of all of the hits showed higher intensity (i.e. gain of ambiguity due to the involvement of multi-genera-hits) (Additional file 1: Fig. S1D). The existing pipelines we tested exhibited the same propensity in detection accuracy (Additional file 1: Fig. S2B). These results point out the inadequacy in the consideration of uniquely mapped reads only and the need for careful handling of multi-genera-hits that causes ambiguity in the contamination source.

To overcome this issue, we designed a scoring scheme for multi-genera-hits (Step VII in Fig. 1A). Based on the overall mapping status of the input reads, multi-genera-hit reads are rigorously penalized when a larger number of uniq-genus-hits are found; however, the penalty is relaxed when uniq-genus-hits are less frequent (Additional file 1: Fig. S3). Overall, our pipeline incorporating this scoring scheme quantifies robust intensities compared to the simple sum of all of the hits (Additional file 1: Fig. S1D). To clarify further, we performed a comparative analysis with the genera detected with over 5% FDR levels in Fig. 1B. The result demonstrated that the loss of accuracy can successfully recover when the weighted multi-genera-hits are considered (Fig. 1C and Additional file 3: Table S1). In addition, our detections of uniq-genus-hits and multi-genera-hits were highly comparable to FastQ screen with Bowtie2, which supports the validity of our mapping strategy tuned with Bowtie2. Interestingly, whereas the local alignment strategies (i.e. PathSeq and FastQ screen) increased the gain of ambiguity, our pipeline reduced it by the scoring scheme.

In this analysis, we observed nine unexpected genera with uniq-genus-hit reads resulting from misalignments for complex reasons (Additional file 3: Table S2). For example, a few reads of *Escherichia coli* were uniquely mapped to Lambdavirus in 3 out of 1,000 runs. To test whether these uniq-genus-hits are rare events, we prepared random reads from our microbe genome database that discarded Lambdavirus genomes and then mapped them to the genera detected in each of the three runs to collect random uniq-genus-hits. After 1,000 runs, in the case of Lambdavirus, the observation of ten unique hits showed almost zero deviation above the mean of the uniq-genus-hits from the mapping of random read sets (p=0.475 with z-score 0.063), implying a chance occurrence of the observed uniq-genus-hits (Additional file 3: Table S2).

Considering these results, we adjusted the proposed method to quantify the microbe abundance at genus-level resolution, and additionally reported species-level quantifications. Evaluation of the significance of the uniq-genus-hits of a genus prior to quantification is critical to avoid false results. For this purpose, instead of adopting the arbitrary criteria used in other methods [9, 14, 16], the proposed pipeline conducts the abovementioned mapping with random read sets to estimate the probability of the occurrence of uniquely mapped reads (Step VIII in Fig. 1A). The genus having significant unique hits is finally quantified by the scoring scheme (Step IX in Fig. 1A).

### Analysis of spike-in contaminants with mesenchymal stem cells

To validate the performance with real-world data, we prepared human periodontal ligament-derived mesenchymal stem cells (hPDL-MSCs) by culturing with and without antibiotic treatments and by adding viable spike-in microbes. We performed DNA-seq, RNA-seq, and ATAC-seq assays with these samples (Table 1). hPDL-MSCs are a promising clinical resource for periodontal regeneration, as studied by our group [30].

**Table 1.**
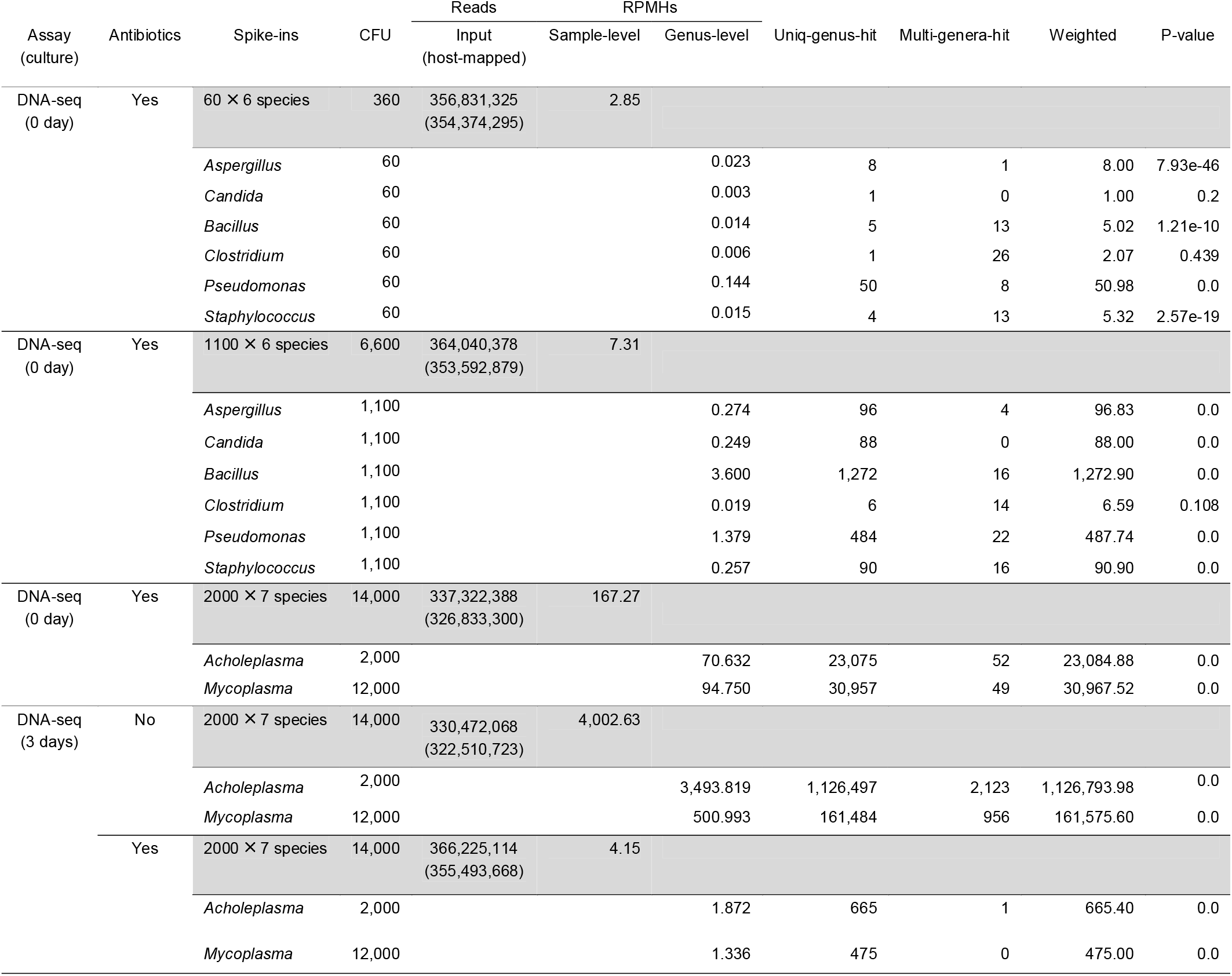
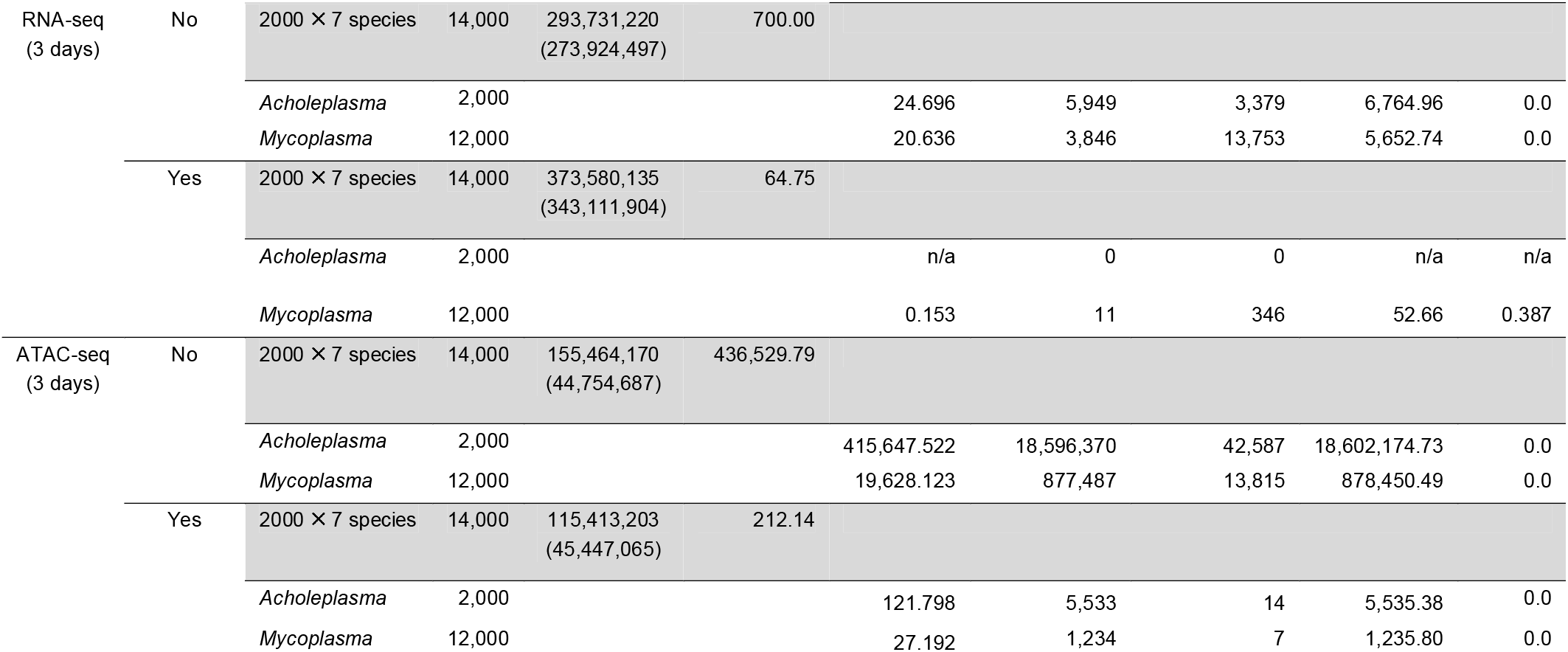
Profiling of spike-in microbes with host-unmapped NGS reads

As shown in Table 1, the spike-in microbes can be quantified with uniq-genus-hits only, decreasing the contribution of weighted multi-genera-hits. In the case of the DNA-seq assay with six spike-in species, we quantified the sample-level RPMHs that were well correlated with the spike-in concentrations (Fig. 1D). At the genus level, we could detect four species at 60 CFU and five species at 1,100 CFU (p<0.001), but failed to detect 60 CFU of *Candida albicans* (p=0.2), as did BWA-align [31] and Taxonomer [17, 32]. By contrast, BWA-mem and NovoAlign found <76 *C*. *albicans* reads with local alignments to low-complexity sequence loci. Of note, the *C. albicans* genome includes a particularly high content of repetitive sequences [33]. These results suggest that the microbial genomic context is one of the factors to determine the detection accuracy particularly in the case of lower contamination degree. In fact, the pipelines increased the detection variability at 60 CFU spike-ins as shown in Fig. 1D; PathSeq with BWA-mem reported a relatively higher concentration and the k-mer matching of Taxonomer broadly reduced the concentrations along with filtering a number of potential host-relevant reads (i.e. 165,777 in Sample1, 85,530 in Sample2, and 84,590 in Sample3).

With regard to antibiotic effects, the DNA-seq assay with 3-day-cultured cells clearly demonstrated that antibiotic supplementation causes a ~1,000-fold decrease in the sample-level RPMH compared with that of cells cultured without antibiotics. In particular, *Acholeplasma* was markedly sensitive to sterilization compared with *Mycoplasma* (Table 1 and Fig. 1E), suggesting the presence of varying drug sensitivities among microbes.

In summary, we concluded that the concentration of spike-in cells can be recovered via our approach. Based on the results of the DNA-seq assays at ~0.1x coverage depth of the host genome with 60 CFU of microbes, we estimated 0.01 RPMH as an approximation of the limit of detection (LOD). That is, one microbial read will exist when 100 million host reads are sequenced. However, LOD verification depends on multiple factors, including microbial genomic context, antibiotic susceptibility, sequencing depth, and sequencing protocol. In this regard, the results of spike-in tests suggest that the ATAC-seq assay offers a remarkable ability to detect contaminants (Fig. 1E) with very few input reads shown in Table 1.

### Detection of prevalent contaminants in public RNA-seq data

To profile the contamination landscape in public data, we downloaded 389 human RNA-seq datasets from ENCODE and Illumina Human BodyMap 2.0 (hereinafter called “IHBM2”), and extracted the potential host-unmapped microbial reads with scattered percentages in the input reads (Additional file 1: Fig. S4A), which amounted to 0.15−18.7% in ENCODE and 0.54−3.0% in IHBM2. Interestingly, the relative level of microbe-mapped reads increased in a sample when the relative level of host-mapped reads decreased (Fig. 2A). Overall, 98% of samples fell within the range of 10^3^−10^5^ RPMHs, forming a reference range for RNA-seq sample-level RPMHs (Fig. 2B).

**Figure 2.**
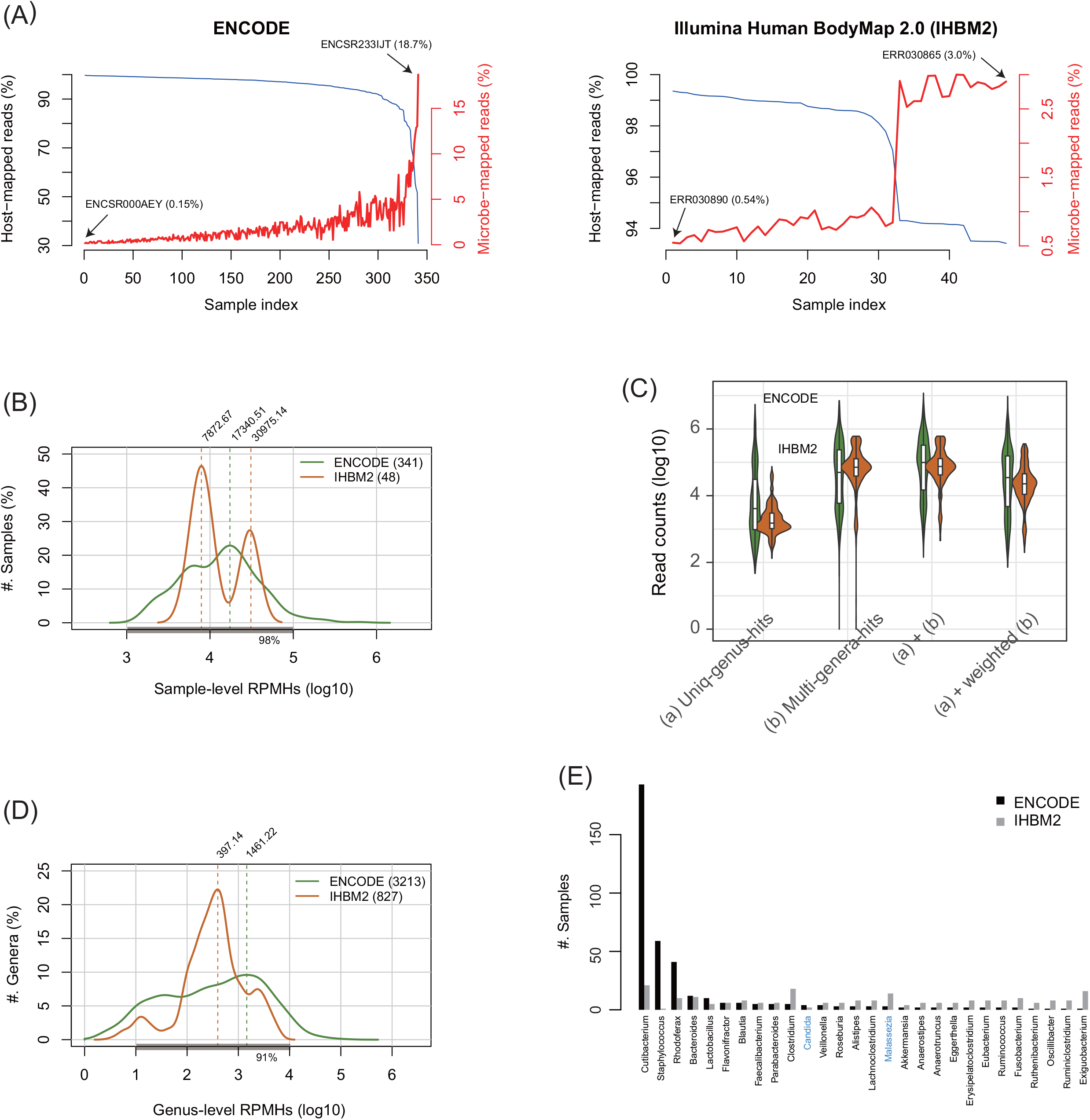
Investigation of 389 public RNA-seq datasets to profile potential contaminants. (A) Distribution of the microbe-mapped reads inversely correlated with that of the host-mapped reads. (B) Distribution of sample-level RPMHs. Of the samples, 98% are within 1,000 to 100,000 RPMHs. (C) Genus-level read counts of 4,040 occurrences of 240 genera across the 389 samples. (D) RPMHs of the 4,040 occurrences, 91% of which are within 10 to 10,000 RPMHs. (E) Twenty-eight genera detected in both ENCODE and Illumina Human BodyMap2.0 (IHBM2) samples; the x-axis labels are colored black for bacteria, blue for fungi, and red for viruses.

At the genus level, we detected 240 genera across the samples (p<0.001). These genera appeared 4,040 times, including widespread multi-genera-hits (Fig. 2C). Using the weighted read counts, we quantified the genus-level RPMHs of the 4,040 occurrences, 91% of which were located within 10 to 10^4^ RPMHs (Fig. 2D). Among the 240 genera, 56 were known contaminants in NGS experiments [12], such as Bacillus, Pseudomonas, and Escherichia (Additional file 1: Fig. S4B). The remainder included 28 genera commonly found in ENCODE and IHBM2 samples (Fig. 2E). In particular, Cutibacterium, including the species *C. acnes* (formerly *Propionibacterium acnes*), which is readily detected on human skin, was the most prevalent, supporting the findings in a previous study [34].

Since the IHBM2 samples exhibited unique patterns, as shown in Fig. 2B and 2D, we next investigated their contamination characteristics by performing cluster analyses. The analysis clearly separated the sequencing libraries and revealed an increased magnitude of contamination in the 16 tissue-mixture samples, likely because producing such samples involved more cell-processing steps (Fig. 3A); this separation led to the bimodal distribution shown in Fig. 2B. To confirm the influence of cell-processing complexity, we further analyzed 22 samples of embryonic stem cells (ESCs) that were sequenced at five time points during culturing on various differentiation media [35]. This analysis revealed three clusters strongly associated with the cell types and time points, and found elevated levels of contamination in the differentiated ESCs (Fig. 3B), suggesting that intricate cell manipulation poses a higher risk of contamination.

**Figure 3.**
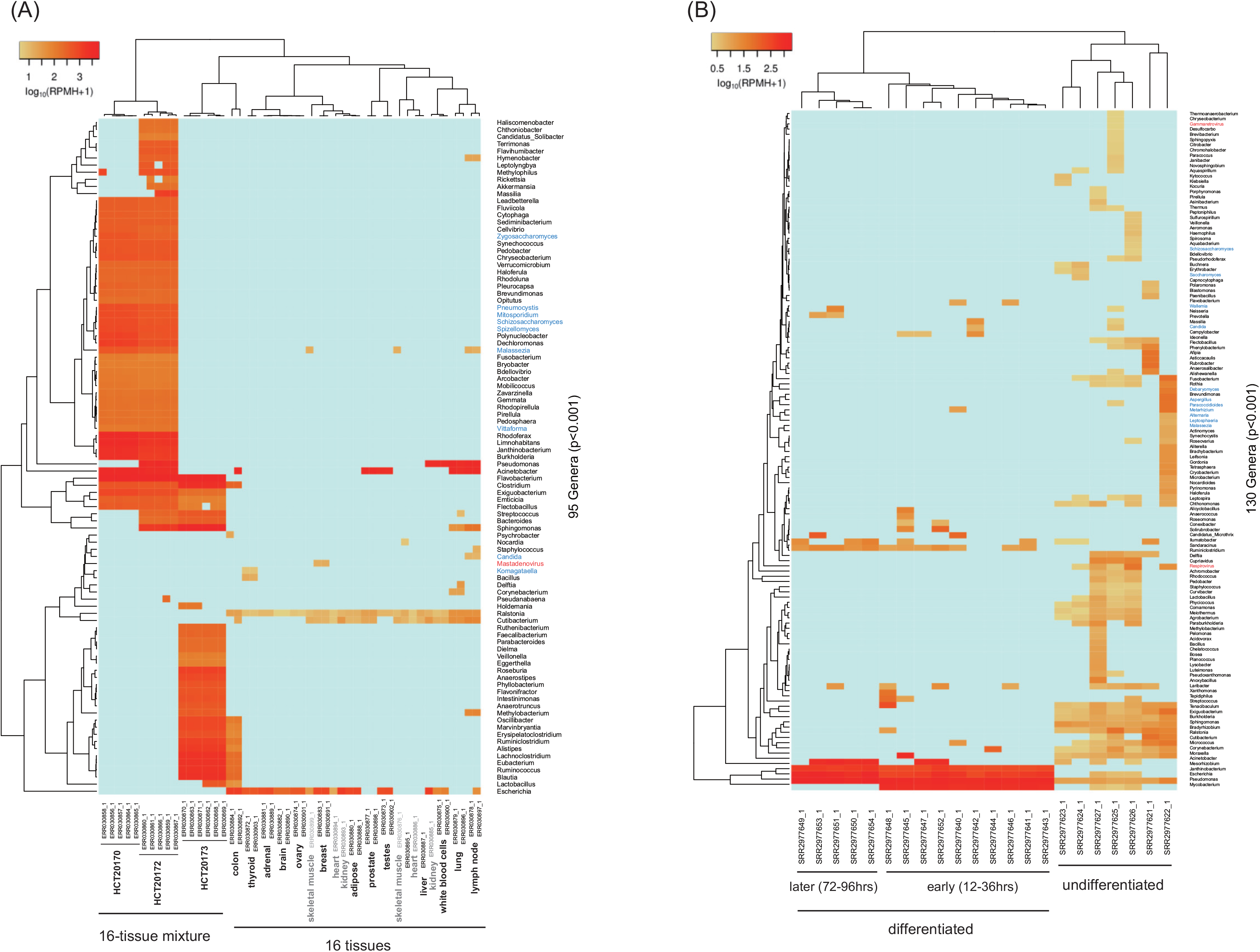
Results of the hierarchical clustering analysis with contamination profiles. (A) Contamination profile of the Illumina Human BodyMap2.0 (IHBM2) samples showing the increased RPMHs in 16 tissue-mixture RNA-seq datasets. (B) Contamination profile of ESCs (SRP067036) showing three clusters associated with differentiation and time points.

Finally, we analyzed host-microbe chimeric reads with paired-end (PE) ENCODE and IHBM2 samples. That is, one end of a PE read was mapped to the host and its counterpart to one or more microbes, and vice versa. The total number of chimeric reads was very low among all of the microbe-mapped reads, implying no considerable influence on the quantification of host gene expression: only 972,812 out of 750,736,667 microbe-mapped PE reads in the ENCODE samples and 93,723 out of 28,622,763 microbe-mapped PE reads in the IHBM2 samples. On the other hand, most of the chimerism existed in host gene bodies that encode ribosome components, transporters, and signaling molecules (Additional file 3: Table S3). The genes were also upregulated in Mycoplasma-infected samples as described below. This finding should be further studied to understand the association between NGS read chimerism and microbial hijacking mechanisms.

### Identifying genes responding to Mycoplasma infection in MSCs

Mycoplasma is notorious for infecting cultured cells and has been frequently detected in public NGS data [8, 9, 36]. Hence, we profiled the genus-level RPMHs of Mycoplasma from the 389 ENCODE and IHBM2 samples as well as from 43 heavily infected samples consisting of seven BL DG-75 samples already known to be infected [9] and 36 lung cancer and stem cell samples. As a result, 110 out of the 432 samples (25.5%) contained at least one Mycoplasma uniq-genus-hit, but only 22 samples (5%) included significant uniq-genus-hits (Fig. 4A). This large discrepancy again suggests the importance of the careful handling of homologous and erroneous NGS reads, which is imperative to infer contaminant prevalence with certainty.

**Figure 4.**
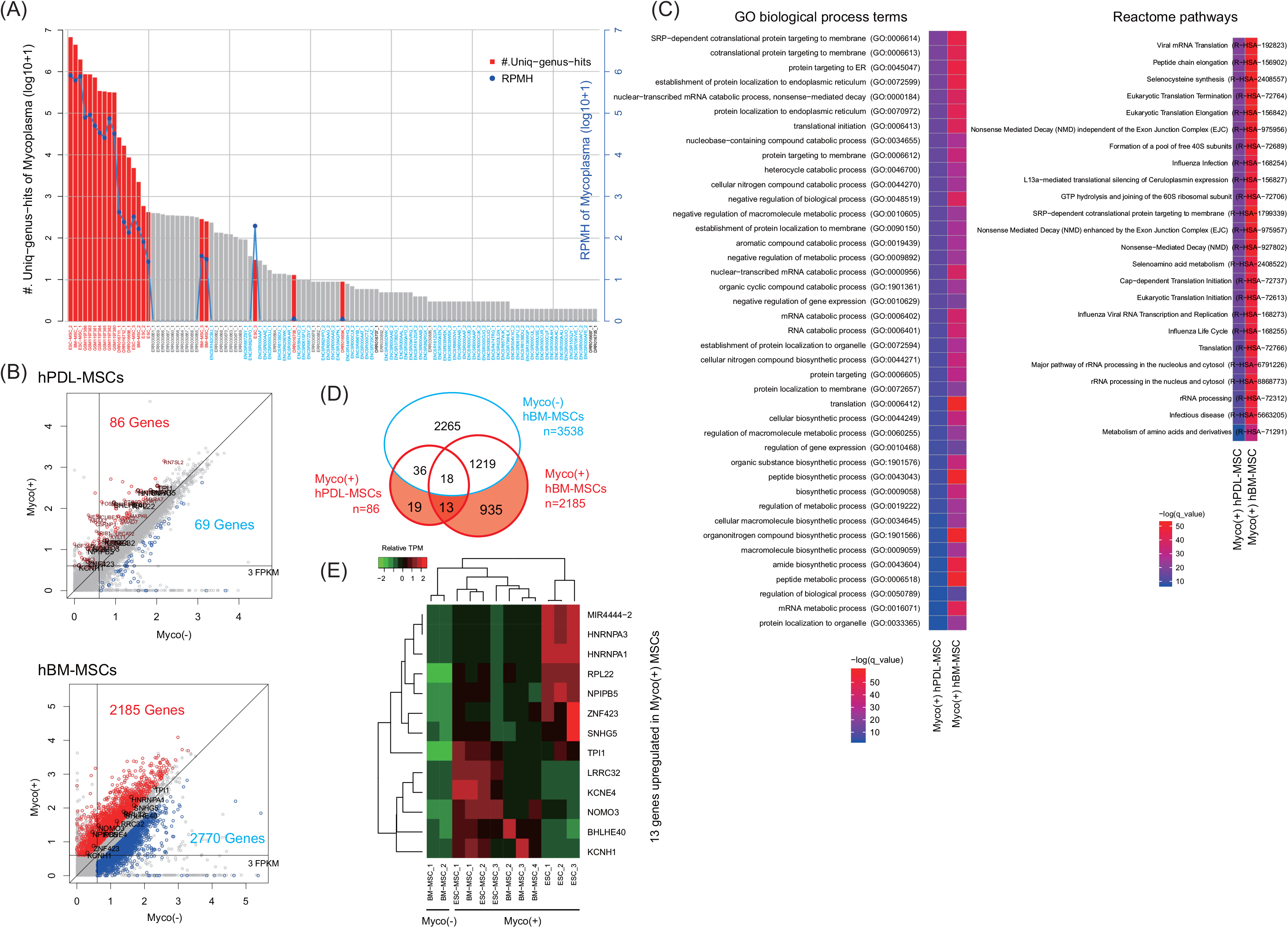
Results of the Mycoplasma prevalence analysis and the functional impacts on host cells. (A) Twenty-two out of 432 public RNA-seq datasets contained significant Mycoplasma-mapped reads (red-colored bar) that were normalized to RPMHs (blue-colored line); the x-axis labels are colored black for DRA001846, gray for IHBM2, blue for ENCODE, and red for Mycoplasma-positive samples. (B) Gene expression correlation plots between Mycoplasma-positive (Myco+) and Mycoplasma-negative (Myco-) MSCs; Myco(+) hPDL-MSCs are Mycoplasma spike-in cells (2000 CFU × 7 species, 3 day cultured without antibiotics), FPKMs were transformed onto the log_10_ scale by adding one, and the black-labeled genes are the 13 genes listed in (D). (C) Highly enriched Gene Ontology terms and Reactome pathways (q-value after Bonferroni correction <0.001). (D) Venn diagram showing unique or shared differentially upregulated genes (DUGs) in MSCs, including 13 out of 967 DUGs unique to Myco(+) MSCs. (E) Expression levels of the 13 genes in Myco(+) ESCs and MSCs; the values are expressed as relative TPM (transcripts per million).

To investigate host gene expression changes during Mycoplasma infection, we identified DEGs between Mycoplasma-positive Myco(+) hPDL-MSCs and uninfected Myco(−) hPDL-MSCs. We performed the same analysis by incorporating the Myco(+) human bone marrow MSCs (hBM-MSCs) used in Fig. 4A and Myco(−) hBM-MSCs (GSE90273). We also sequenced and identified DEGs from Myco(−) hBM-MSCs as a control. Of note, although decreases in gene expression should also be studied, we focused on the differentially upregulated genes (DUGs) in the Myco(+) samples to enable clear interpretations. We identified 86 and 2,185 DUGs in Myco(+) hPDL-MSCs and in Myco(+) hBM-MSCs, respectively (Fig. 4B), 31 of which existed in both classes of MSCs. Although the DUGs are broadly involved in RNA processing, the genes are significantly enriched in cotranslational protein transport processes and with pathways involved in infection responses (Fig. 4C). None of these enrichments were observed among the 3,538 DEGs in Myco(−) hBM-MSCs (Additional file 1: Fig. S5). Among the 967 DUGs identified in Myco(+) MSCs, we ultimately retrieved 13 genes that are specifically upregulated in Myco(+) hPDL-MSCs and hBM-MSCs (Fig. 4D).

These results imply that the Mycoplasma in the MSCs addressed here utilizes host protein biosynthesis machinery related to the ER-associated degradation (ERAD) pathway, a well-known microbial entry point [37, 38]. Moreover, one can infer that the abnormal increase in the expression levels of the 13 DUG RNAs is a candidate diagnostic marker for infection. Indeed, the DUGs were also upregulated either in Myco(+) ESCs or other Myco(+) MSCs (Fig. 4E).

### Inference of the functional impact of multiple contaminants

As shown in Fig. 5A, a few genes among the 967 DUGs in the Myco(+) MSCs were upregulated in Myco(+) DG-75 samples, which suggests a different type of response in lymphoma. We investigated the correspondence between gene expression levels and Mycoplasma concentrations in the samples and identified genes potentially associated with the infection (Additional file 1: Fig. S6A); however, significant GO terms were not detected, which is consistent with the findings of a previous report [9]. Remarkably, the DG-75 samples were heavily contaminated with multiple microbes (Fig 5B), and the gene expression levels exhibited diverse correlation patterns with the concentrations of other microbes (Additional file 1: Fig. S6B), implying a profound influence of co-contaminants on phenotypes.

**Figure 5.**
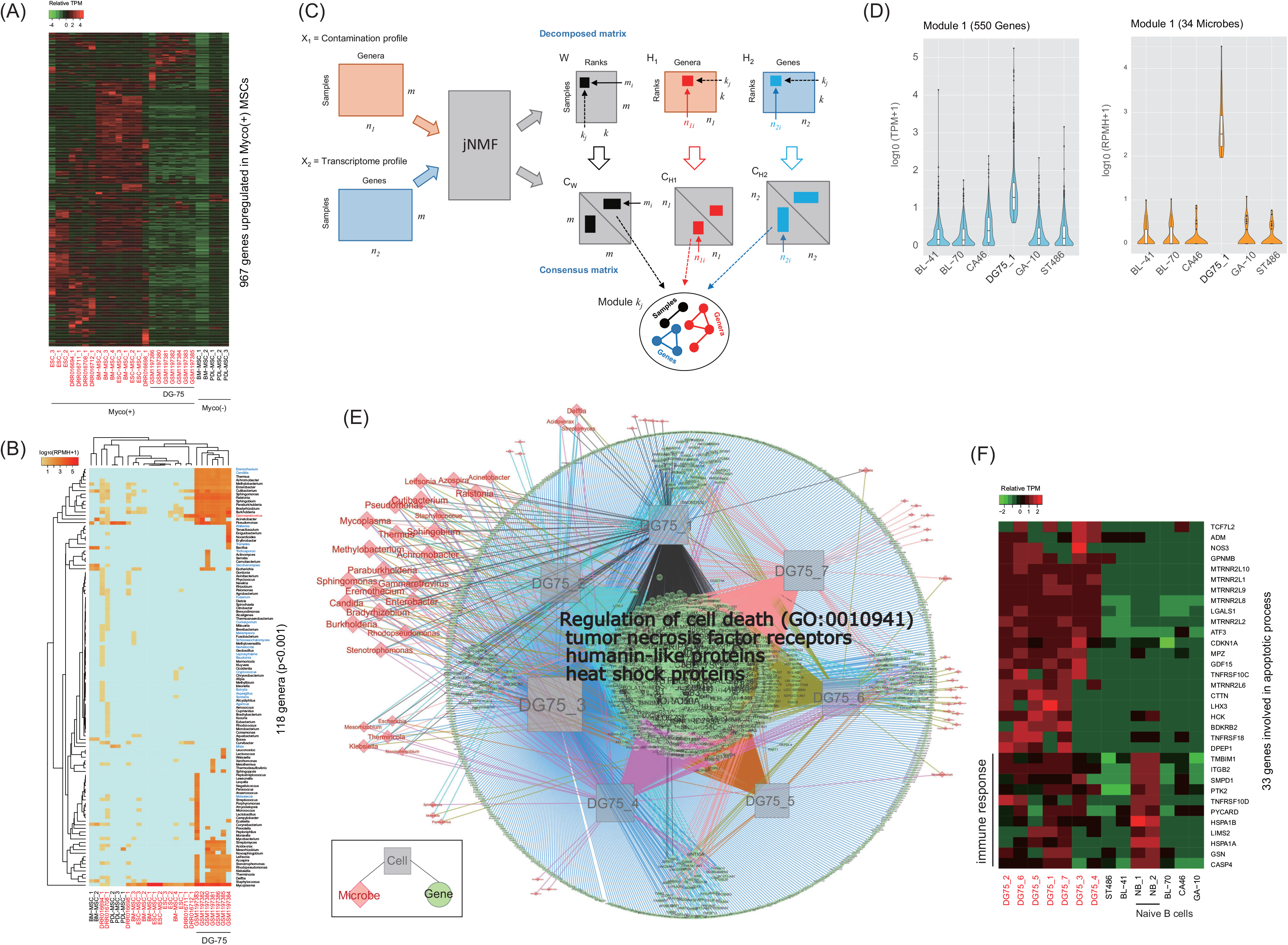
Inference of DUGs associated with multiple contaminants in Myco(+) DG75 samples. (A) Expression profile of 967 DUGs unique to Myco(+) MSCs. (B) Contamination profile with MSC, ESC, and DG-75 samples; the x-axis labels are colored black for Myco(−) and red for Myco(+). (C) Schematic representation of module identification from two input profiles by the jNMF algorithm. (D) An example showing the module that captured genes and contaminants co-elevated in a DG-75 sample. (E) Network representation of the association between genes and contaminants co-elevated in the seven DG-75 samples; GO:0010941 is the enriched GO term in the genes found in at least four DG-75 samples (p=3.76e-3). (F) Expression profiles of the 33 genes involved in the biological process “regulation of cell death”; DG75_1 (GSM1197380), DG75_2 (GSM1197385), DG75_3 (GSM1197386), DG75_4 (GSM1197381), DG75_5 (GSM1197382), DG75_6 (GSM1197383), DG75_7 (GSM1197384), NB_1 (GSM2225743), and NB_2 (GSM2225744).

To facilitate the inference of the impact of multiple contaminants, we employed a joint non-negative matrix factorization (jNMF) algorithm [39, 40] that modulates multiple genes and contaminants associated in a set of samples (Fig. 5C). We first prepared seven input datasets, each of which contained five Myco(−) BL cell lines and one of the seven Myco(+) DG-75 samples. After preparing contamination and transcriptome profiles for each dataset, we repeatedly ran the jNMF algorithm by setting a series of parameters for testing the clustering stability (Additional file 1: Fig. S7). In the case of DG75_1 (GSM1197380), the jNMF algorithm retrieved the module that specifically includes elements co-elevated in the dataset, i.e., 550 genes and 34 contaminants, including Mycoplasma (Fig. 5D). By gathering this type of module from all of the results of the seven input datasets, we could build a network modeling the connectivity between the upregulated genes and microbe concentrations in the DG-75 samples (Fig. 5E).

The network consisted of 4,322 edges connecting 2,289 genes, 68 microbes, and seven samples. Of these genes, 259 genes were common to least four DG-75 samples, and the biological process “*regulation of cell death*” (GO:0010941) was significantly enriched in a subset of them (p=3.76e-3). This subset (33 genes) included tumor necrosis factor receptors, which paradoxically play pro-tumorigenic or pro-apoptotic functions [41], and humanin-like proteins, which potentially produce mitochondria-derived peptides that inhibit apoptosis [42]. Some of the genes were also highly expressed in normal B cells, where they are likely involved in activating immune responses. The Myco(−) BL cell lines exhibited repression of these apoptosis-related genes (Fig. 5F), which implies that the effect is not specific to cancerous cell types.

These results suggest that the severely contaminated DG-75 samples resisted contamination by multiple microbes via inflammation pathways and survived by inhibiting apoptotic pathways via mitochondria-related mechanisms or via the inhibitory effect of Mycoplasma on apoptosis [36]. Collectively, we concluded that jNMF facilitates the inference of how phenotypes (i.e., gene expression in this case) have been affected by the complex activities of co-contaminants.

## Discussion

We sought to assess the feasibility of NGS-based contaminant detection and to improve its certainty by conducting microbe-spike-in experiments and by analyzing public data. For profiling microbial contamination, the use of metagenomics approaches that depend on phylogenetic markers or de novo assembly seem to offer little benefit, because the sterilization of microbes and sequencing library preparation from host cell DNA lead to dilution and degradation of microbe-derived nucleic acids [13, 14]. Furthermore, since microbial communities can contaminate host cells, a comprehensive catalog of microbial genomes must be considered to avoid false inferences. Preliminarily, we detected phiX174 in 77 out of 341 ENCODE samples with the numbers of mapped reads ranging from 177 (ENCSR000AEG) to 7,031,626 (ENCSR000AAL). Surprisingly, fewer than six reads in a sample were the uniq-genus-hits of phiX174, and the remainder were multi-genera-hits for phylogenetic neighbor bacteriophages [24, 43, 44]. This situation, which makes it difficult to identify the true species, may occur frequently, as the uniquely mapped and multi-mapped reads in the public datasets exhibited a broad range of intensities (Fig. 2C).

We here developed a straightforward approach that uses a large-scale genome database and exploits multi-mapped reads that were discarded in previous studies. Although our method successfully detected the origins of microbes from the simulated reads of random mixtures, the detection certainty was still imperfect, particularly at species level resolution. To overcome this issue, we attempted to estimate whether unique microbe-mapped reads are likely observed by chance. We found that 80% of the 110 public RNA-seq samples in which uniq-genus-hits of Mycoplasma were detected resulted from random occurrences, and 5% of 432 RNA-seq samples were most likely infected with Mycoplasma. Moreover, we estimated 10^3^−10^5^ sample-level RPMHs consisting of 10−10^4^ genus-level RPMHs, consistent with previous reports; however, these results illustrated more dispersion than expected. Of note, it is possible that these RPMH estimations are limited to the samples used here, as microbes are highly sensitive to environmental conditions due to distinct genomic context, growth rate, antibiotic susceptibility, and invasion mechanism, and RPMH distributions depend greatly on the sample sets analyzed.

As shown by the results of the spike-in analyses, even though the experimental conditions were identical, the profiles differed between the DNA-seq, RNA-seq, and ATAC-seq assays. Remarkably, RNA-seq profiling tended to include more diverse microbes. This tendency may be attributed to the relatively complex sample handling required, which leads to a higher risk of contamination. Indeed, elaborate cell manipulations, such as tissue mixture and induction of cell differentiation, result in increased contamination diversity and intensity. On the other hand, because most prokaryotes have histone-free supercoiled nucleoids [45], ATAC-seq is superior for microbe detection with very low numbers of input reads. This suggests that the ratio of microbe-to-human DNA accessibility is useful to the NGS-based microbial contaminant detection more than the ratios of the genome and transcriptome sizes. This aspect of our work should be explored in more detail in future studies.

By analyzing public NGS samples, we found that microbes from the genus *Cutibacterium* are widespread contaminants, which is thought to arise naturally [12]. In addition to known contaminants, our microbe catalog suggests that the major sources of contamination are laboratory reagents and experimental environments. Importantly, any microbial contamination can trigger phenotypic changes in the host cells; however, the response pathways are diverse and unclear. For example, the genes aberrantly expressed during Mycoplasma infection differed greatly between MSCs and cancer cells. Therefore, as an approach to systematically infer the effects of contamination, we used network analysis with jNMF. This approach revealed that host-contaminant interactions alter the molecular landscape, and such alterations could result in erroneous experimental conclusions.

## Conclusions

The findings in this study reinforce our appreciation of the extreme importance of precisely determining the origins and functional impacts of contamination to ensure quality research. In conclusion, NGS-based contaminant detection supported by efficient informatics approaches offers a promising opportunity to comprehensively profile contamination landscapes.

## Methods

### Step-by-step procedure of the proposed pipeline

The proposed pipeline shown in Fig. 1A consists of step-by-step operations detailed below.

Step I (Quality control): Trimmomatic [46], with the option “ILLUMINACLIP:*adapter_file*:2:30:10 LEADING:20 TRAILING:20 MINLEN:36”, assesses the quality of the input NGS reads by removing adapters and trimming reads.

Step II (Mapping to host-reference genome): HISAT2 [47] coupled with Bowtie2 [27] with the option “-k 1” aligns the quality-controlled reads to a host reference genome.

Step III (Removing host-relevant reads): To remove any potential host reads, Bowtie2 with “--sensitive” and via BLASTn with the options “-evalue 0.001 -perc_identity 80 -max_target_seqs 1” sequentially align the unmapped reads again to alternative host genomic and transcriptomic sequences.

Step IV (Making low-complexity sequences): The host-unmapped reads that still remain are candidate contaminant-origin reads. To reduce false discovery, TANTAN [48] masks the low-complexity sequences in the host-unmapped reads.

Step V (Mapping to a microbe genome): Bowtie2, with the option “--sensitive”, aligns the masked sequences to one set of bacterial, viral, or fungal genomes of species belonging to the same genus. This step is independently repeated with each of the 2,289 genera.

Step VI (Categorizing read-mapping status): A mapped read is categorized as either a ‘uniq-genus-hit’ (i.e. uniquely mapped to a specific genus) or a ‘multi-genera-hit’ (i.e. repeatedly mapped to multiple genera). The statistics is gathered from the mapping results, which includes the total number of microbe-mapped reads (i.e. sum of ‘uniq-genus-hit’ and ‘multi-genera-hit’) and the total number of host-mapped reads.

Step VII (Defining a shape of scoring function): The total number of microbe-mapped reads (*n*) and the number of genera of each ‘multi-genera-hit’ read (*T_i_*) define an exponential function for weighting the ‘multi-genera-hit’ reads. That is, a score *S_i_* for the read *i* that was mapped to *T_i_* different genera (or a single genus) is given by

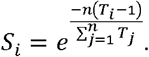

Thus, a read uniquely mapped to a genus is counted as 1.0, whereas a read mapped to multiple genera is penalized by the exponential function.

Step VIII (Testing statistical significance of unique hits): To test the chance occurrence of the ‘uniq-genus-hit’ reads that were mapped to specific microbes, the pipeline first randomly samples *n* reads (i.e. the total number of microbe-mapped reads) from the microbe genomes that discard the observed microbial genomes. Next, the pipeline aligns the random reads to the observed microbial genomes and counts the uniquely mapped reads. This procedure is repeated ten times to prepare an ensemble of random numbers of unique reads for each observed genus. The numbers for a genus are converted into z-scores, and the null hypothesis that no difference exists between the observation and the mean of its ensemble is tested, resulting in a p-value.

Step IX (C alculating RPMHs): For sample-level quantification, a normalized RPMHs core (read per million host-mapped reads) is calculated as RPMH =*n*/*m* × 10^6^, where *n* and *m* are the total number of microbe-mapped reads and the total number of host-mapped reads in a given input dataset, respectively. For genus-level quantification, the RPMH of a genus *G* is calculated by

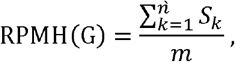

where *ǹ* is the total number of reads uniquely or repeatedly mapped to *G*.

### Preparation of random microbial reads for reversion

Ten species belonging to distinct genera were randomly selected and prepared 1,000 100-base pair (bp) DNA fragments from the genome of a selected species were prepared. A run of the reversion test uses the 10,000 reads (1,000 reads × 10 species) and calculates the false discovery rate (FDR) for each species; that is, TN / (TN+TP), where TP (true positive) is the number of reads mapped to their origin and TN (true negative) is the number of reads mapped to others. If the method works perfectly, the species tested will be detected with 1,000 uniquely mapped reads (see Additional file 2).

### Cell collection and culture

Human bone marrow-derived MSCs (hBM-MSCs) were purchased from Lonza (Lonza, Walkersville, MD, USA), and periodontal ligament-derived MSCs (hPDL-MSCs) were prepared as previously described [49]. Briefly, periodontal ligament (PDL) tissue samples separated from the middle third of a patient’s wisdom tooth were digested with collagenase (Collagenase NB 6 GMP Grade from *Clostridium histolyticum*; Serva, Heidelberg, Germany)/dispase (Godo Shusei Co., Tokyo, Japan), and single-cell suspensions were passed through a 70 μm cell strainer (Falcon, Franklin Lakes, N.J., USA). The collected cells were incubated in a culture plate (Falcon T-25 flask, Primaria; BD Biosciences, San Jose, CA, USA) in complete medium: α-MEM (Sigma-Aldrich, St. Louis, MO, USA) containing 10% fetal bovine serum (Gibco; Thermo Fisher Scientific, Waltham, MA, USA), 2 mM L-glutamine (Sigma-Aldrich, St. Louis, MO, USA), and 82.1 μg/ml L-ascorbic acid phosphate magnesium salt n-hydrate (Wako Junyaku, Tokyo, Japan) with the antibiotics gentamicin (40 μg/ml, GENTCIN; Schering-Plough, Osaka, Japan) and amphotericin B (0.25 μg/m, FUNGIZONE; Bristol-Myers Squibb, Tokyo, Japan). After three passages for expansion in T-225 flasks, the cells were preserved in freezing media (STEM-CELLBANKER GMP grade; Nihon Zenyaku Kogyo, Fukushima, Japan) and stored in liquid nitrogen.

### Spike-in test of microbes with human PDL-MSCs

The frozen cells were rapidly thawed with gentle shaking in a water bath at 37 °C. Next, the cells were spiked and cultured in complete medium with and without antibiotics (40 μg/ml gentamicin and 0.25 μg/m amphotericin B). Then, 2 × 10^5^ cells were spiked with either Bioball^®^ (BioMérieux, France) or seven species of *Mycoplasma* (Additional file 3: Table S4), 60 or 1,100 colony forming units (CFU) of each Bioball, or 2,000 CFU of each *Mycoplasma* species. Genomic DNA was isolated 0 or 3 days after the spike-in using a NucleoSpin Blood Kit (Macherery-Nagel Inc., Easton, PA, USA), and total RNA was isolated using a NucleoSpin RNA kit (Macherery-Nagel Inc., Easton).

### Sequencing of DNA and RNA libraries

DNA-seq libraries were prepared using 100 ng DNA and the Illumina TruSeq Nano Kit, following the manufacturer’s instructions. RNA-seq libraries were prepared using 200 ng total RNA and the SureSelect Strand-Specific RNA Reagent Kit (Agilent Technologies, Santa Clara, CA, USA), following the manufacturer’s instructions. ATAC-seq libraries were prepared using 50,000 cells, according to a published protocol [50]. Sequencing of 36 bp single ends of the RNA libraries from mycoplasma-free hPDL-MSCs (three biological replicates) and hBM-MSCs (three biological replicates) was performed with an Illumina HiSeq2500 system. Sequencing of the 100 bp paired ends of the libraries of hPDL-MSCs with microbe spike-in was conducted with an Illumina HiSeq3000 system.

### Implementation of joint non-negative matrix factorization

Joint non-negative matrix factorization (jNMF) has been successfully applied for the detection of so-called “modules” in multiple genomic data [40, 51, 52]. Briefly, given *N* multiple non-negative data matrices 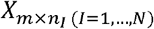, jNMF decomposes the input matrices into a common basis matrix *W*_*m*×*k*_ and a set of coefficient matrices 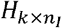by minimizing a squared Euclidean error function formulated as

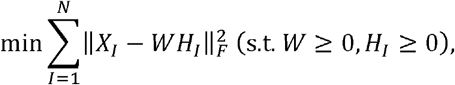

where *k* is the factorization rank and *F* is the Frobenius norm. To optimize this objective function, a multiplicative update procedure was performed by starting with randomized values for *w* and *H*_*1*_, which is well described in many publications [40, 51, 53]. In single trial, the update procedure was repeated *R* times, and the trial was restarted *T* times. During the trials, consensus matrices *c*_*m*×*m*_ and 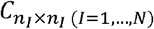 were built to calculate the co-clustering probabilities of all of the input elements, i.e., the cophenetic correlation coefficient values [39]. For example, if the maximal value of the *j*th factorization rank coincides with the *i*th element in *W*_*m*×*k*_, all of the elements in *m* having >0.8 with the *i*th element in *c*_*m*×*m*_ were modulated. In this study, *N* = 2 (i.e., contamination profile and expression profile) and *m* = 6 (i.e., five Myco(−) samples and one Myco(+) sample) were used. Thus, *m*, *n*_1_, and *n*_2_ represent cells, contaminants, and genes, respectively. The parameters *T* = 100,*R* = 5000, and *k* = 3 were set after testing the clustering stabilities with the combinations of *T* = (10,50,100), *R* = (1000,2000,5000), and *k* = (2,3,4,5) by calculating the cophenetic correlation coefficient values [39]. The input profiles retaining elements with >3 TPM and >1 RPMH were converted to the log_10_ scale by adding one.

### Preparation of public datasets

The human reference genome (hg38) was downloaded from the UCSC genome browser [54] and alternative sequences of the reference genome were downloaded from the NCBI BLAST DB [55]. To build the microbial genome database, the complete genomes of bacteria, viruses, and fungi were obtained from the NCBI RefSeq [56], consisting of 11,360 species from 2,289 genera. Raw RNA-seq datasets (341) were downloaded from the ENCODE project [57] and additional raw RNA-seq datasets were downloaded from NCBI’s GEO and SRA, including 48 Illumina Human BodyMap 2.0 (GSE30611), 22 ESCs (SRP067036), seven Burkitt’s lymphoma (BL) DG-75 cell lines (GSE49321), 26 lung cancer cell lines (DRA001846), and ten stem cells (PRJNA277616). The RNA-seq data for the EBV-negative BL cell lines (BL-41, BL-70, CA46, GA-10, and ST486) were obtained from the CCLE [58].

### Bioinformatics analysis

To analyze the RNA-seq data, the HISAT2-Bowtie2 pipeline and the Cufflinks package [47, 59] were used with hg38 and RefSeq gene annotation. After retrieving genes with >3 FPKMs in at least one sample, Cuffmerge and Cuffdiff were performed to detect differentially expressed genes (DEGs) satisfying a q-value cutoff <0.05 (Benjamini-Hochberg correction p-value) and a >2.0 fold-change (fc) cutoff. To analyze the RPMH clusters, R language function hclust was used. The Euclidean distances among the RPMHs were adjusted by quantile normalization and mean centering, and the hierarchical average linkage method was used to group genera. To analyze the enrichment of Gene Ontology (GO) terms and pathways, the GOC web tool [60] was used with the “*GO biological process complete*” and “*Reactome pathways*” datasets by selecting the option “*Bonferroni correction for multiple testing*”.

NovoAlign (V.3.08) was downloaded from the Novocraft [61] and Taxonomer was performed on the Taxonomer website [32]. The network data were visualized by using software Cytoscape (V.3.5.1). PathSeq [18], FastQ Screen [28], and DecontaMiner [29] were installed with their reference databases. Because FastQ Screen accepts limited number of genomes, the input reads were mapped to 10 specific genomes only. Detailed information on the existing pipelines can be found from Additional file 2. To calculate the sample-level RPMHs in Fig. 1D, the existing pipelines were used to analyze the host-unmapped reads of our pipeline, and the total number of microbe-mapped reads was divided by the total number of host-mapped reads from our pipeline. As the total number of microbe-mapped reads, for Taxonomer, the numbers of ambiguous, bacterial, fungal, phage, phix, and viral bins in the output file were summed up. For DecontaMiner, the total counts of ‘TOTAL_READS’ in the output file were collected. For PathSeq, the read count of the column ‘read’ when the column ‘type’ is ‘root’ in the output file was collected.

## Availability of data and materials

All data generated or analyzed during this study are included in this published article and its supplementary information files. The DNA-seq, RNA-seq, and ATAC-seq data have been deposited in the NCBI Sequence Read Archive (SRP161443) [62]. The source code of jNMF and the dataset for Fig. 1B have been deposited in GitHub [63]. The online version of the proposed pipeline is available at our web site [64]. The scripts and materials are available from the corresponding author on request.

## Additional files

### Additional file 1

**Figure S1.** Results of the reversion test employing different parameters for Bowtie2. Using the simulated read sets created in Fig. 1B, Bowtie2 was performed with the parameters “--very-sensitive” (A), “--fast” (B), and “--very-fast” (C). (D) Distribution of the reverted reads of 5,709 species at genus-level resolution (“--sensitive” parameter).

**Figure S2.** Results of the reversion test in the three existing pipelines. (A) FDR distributions at genus-level resolution. (B) Distribution of the reverted reads of 5,709 species at genus-level resolution. Additional file2 details how these values were calculated.

**Figure S3.** Examples of the scoring function used to weight multi-genera-hit reads. The slope of the exponential function is defined by the overall mapping status of the input reads incorporated into *M* (the total number of microbe-mapped reads) and *N* (the total number of unique or multiple hits of all microbe-mapped reads). For instance, a read of ENCSR000AAR that mapped to ten distinct genera (*T*=10) is counted as 0.4.

**Figure S4.** Profiling contamination prevalence in public RNA-seq datasets. (A) Distributions of the fractions of microbe-mapped reads in the total input reads of ENCODE and IHBM2 (Illumina Human BodyMap 2.0). (B) Frequencies of 240 microbial genera detected as significant contaminants in the samples. The gray-colored bars represent known contaminants reported in Salter, et al., 2014 [12]. Microbes labeled in black-, blue-, and red correspond to bacterium, fungus, and virus, respectively.

**Figure S5.** Results of the enrichment analysis of GO biological process terms with DEGs found in Myco(−) hBM-MSC BM1 and BM2. In BM1 and BM2, 2237 DUGs (differentially upregulated genes) and 1301 DUGs were identified, respectively. The heatmap showed over-enriched GO terms in both BM1 and BM2. The enrichment analysis of the reactome showed no significant enrichments (q-value <0.001). DUG_BM1: differentially-upregulated genes in Myco(−) hBM-MSCs that were sequenced in this study, DUG_BM2: differentially-upregulated genes in Myco(−) hBM-MSCs that are publicly available (GSE90273). The q-value is the Bonferroni-corrected p-value for multiple testing.

**Figure S6.** Correlation analysis of gene expression with Mycoplasma concentration in Myco(+) DG-75 samples (GSE49321). (A) Genes that exhibited positively (94) or negatively (195) correlated expression patterns with Mycoplasma RPMHs among the seven samples (>0.8 or < −0.8 in Pearson’s correlation coefficient); gene expression levels are relative TPM values. (B) Distribution of correlation coefficient values of TPM values with multiple contaminant RPMHs.

**Figure S7.** jNMF example with 5 Myco(−) BL cell lines and Myco(+) DG-75_3 (GSM1197386). (A) Contamination and gene expression profiles addressed by jNMF. (B) Distributions of Cophenetic correlation coefficient (Cophenetic CC) values and RSSs (residual sum of squares) in different *k*-ranks (=2,3,4,5). Cophenetic CC values show the clustering stability and RSS represents the difference between the matrices (A) and reconstructed matrices by jNMF; each box at a *k* -rank includes the results from 9 jNMF runs with different parameters; (1) results with the matrices (A); (2) results with randomized matrices of (A). At rank *k*>3, (1) and (2) became indistinguishable via the Cophenetic CC, suggesting that the choice of *k*=3 was reasonable. Other parameter sets were not influenced in (1). (C) The common basis matrix W and consensus matrices estimated by jNMF with the parameters *k*=3, T=100 and R=5000. (D) jNMF modules found in (C); K2 in (C) corresponds to Module 1. (PDF 4,426 kb)

### Additional file 2

Concept of the reversion test and its procedure for existing pipelines (PDF 117kb).

### Additional file 3

**Table S1.** List of 11 genera of 291 reversion-test runs showing over 5% FDR level.

**Table S2.** List of genera accidentally found in the reversion tests with different Bowtie2 parameters.

**Table S3.** Host-microbe chimeric reads overlapped with intergenic or host gene body regions.

**Table S4.** List of spike-in microbes. (XLSX 135kb)

