## Supplementary figures and images for "A systematic NGS-based approach for contaminant detection and functional inference"

### Additional file 1

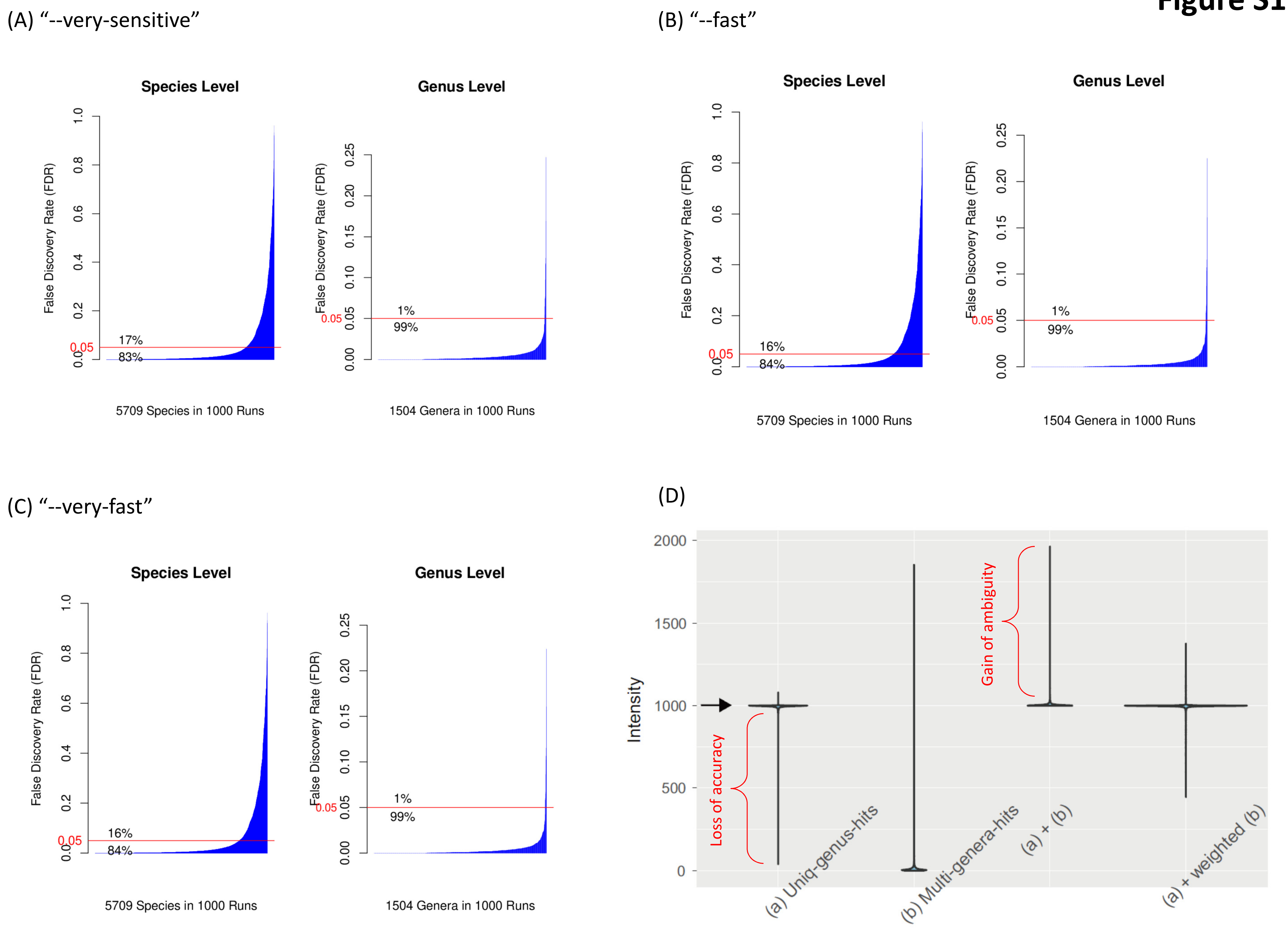

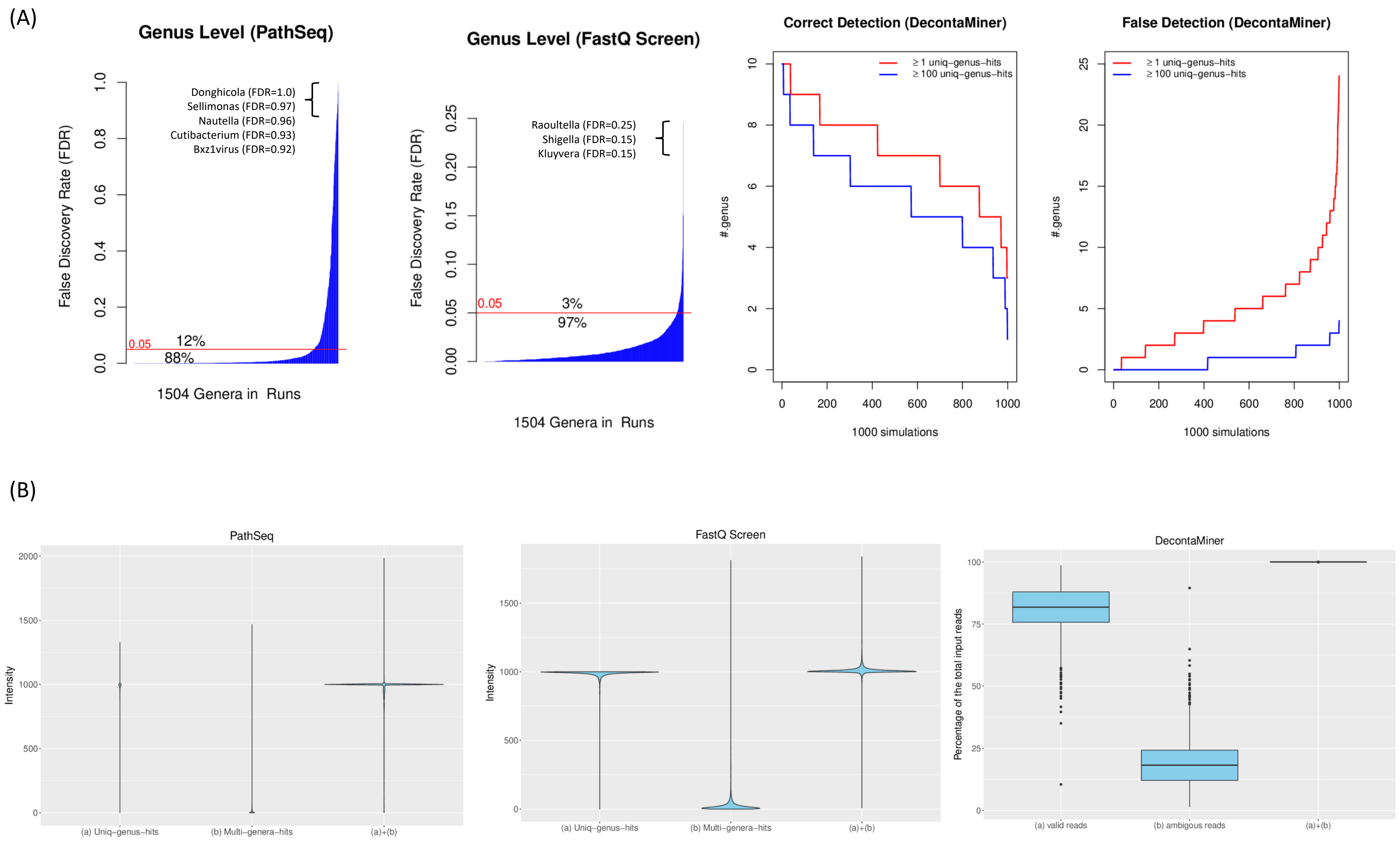

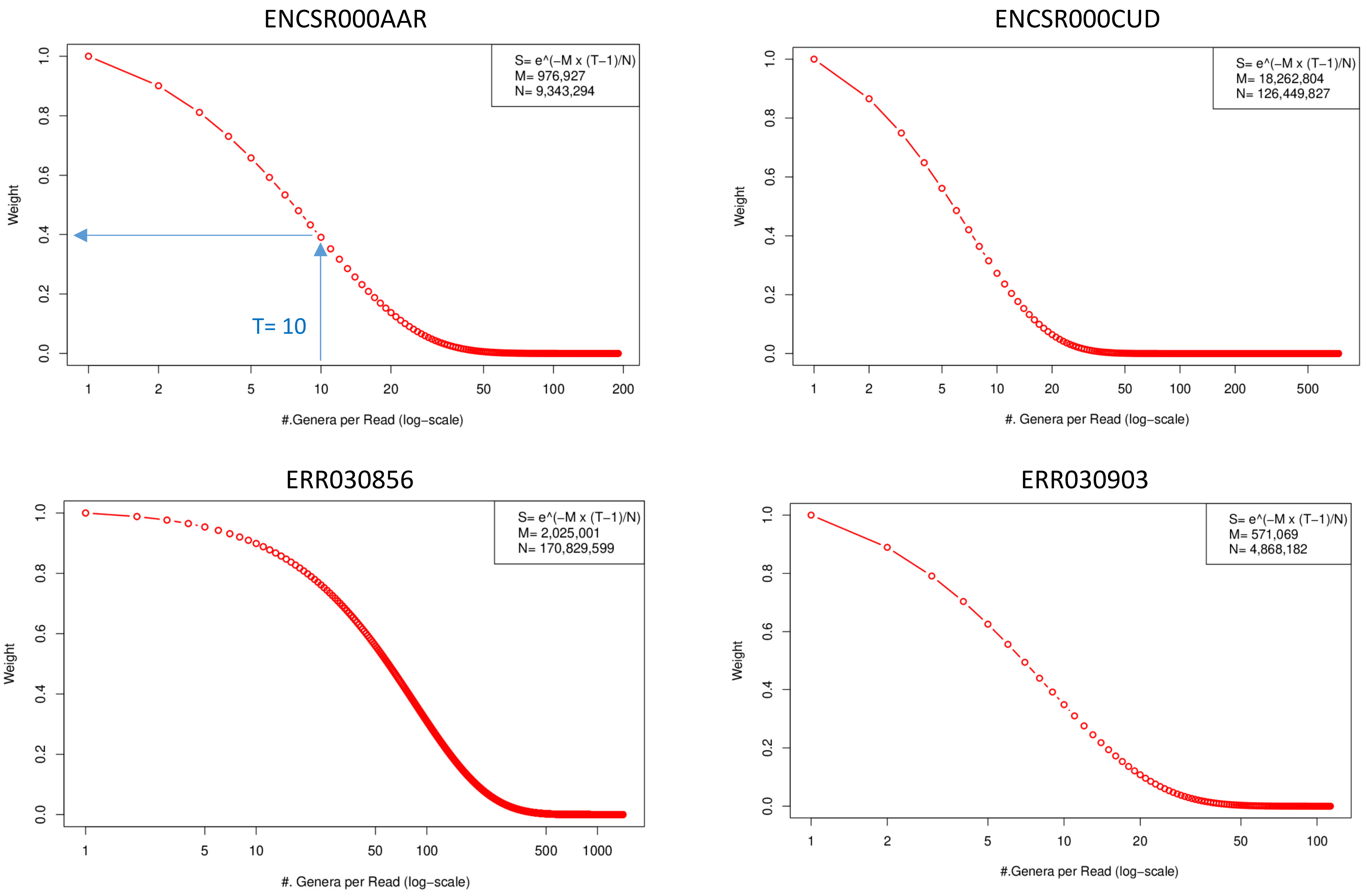

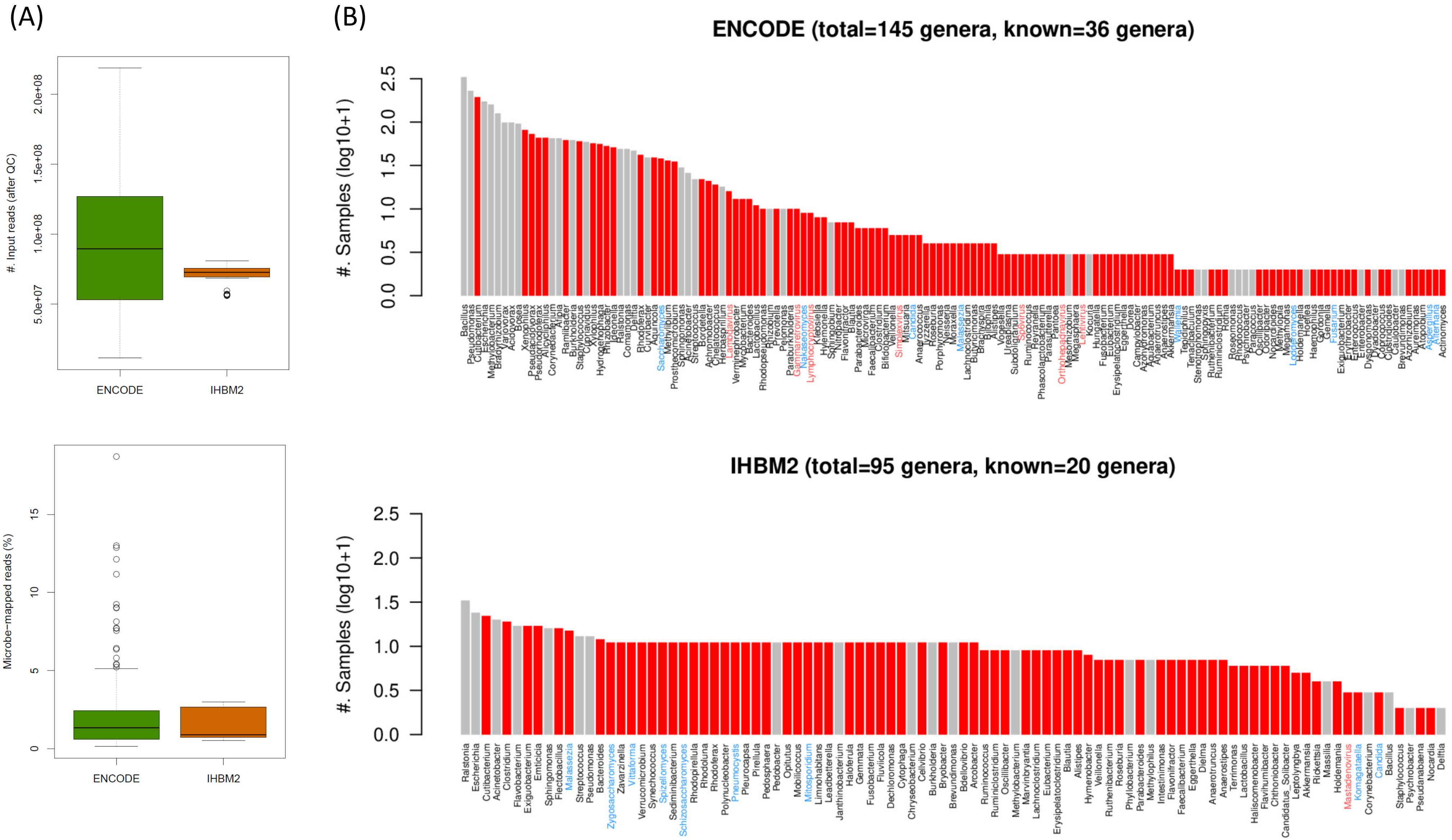

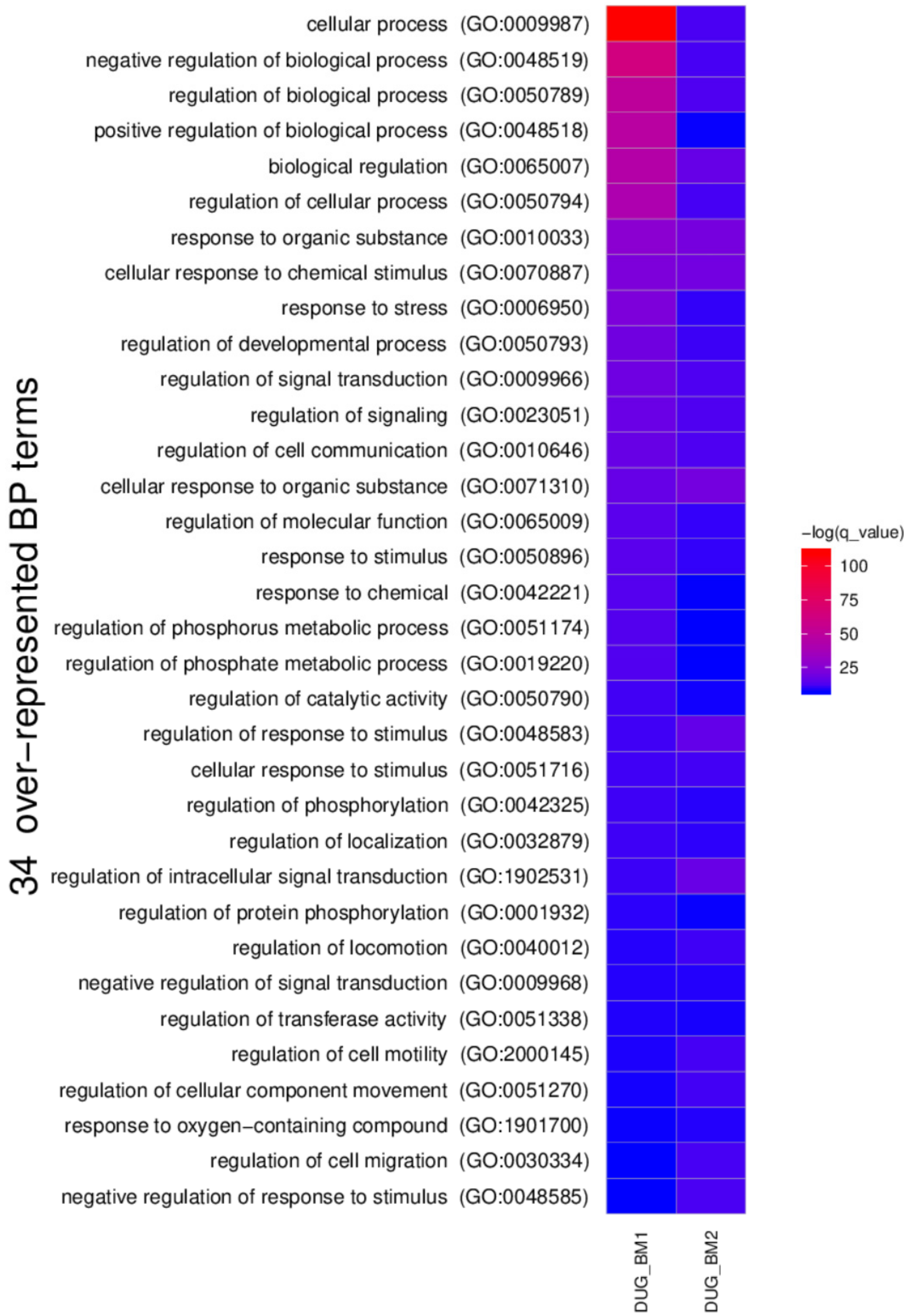

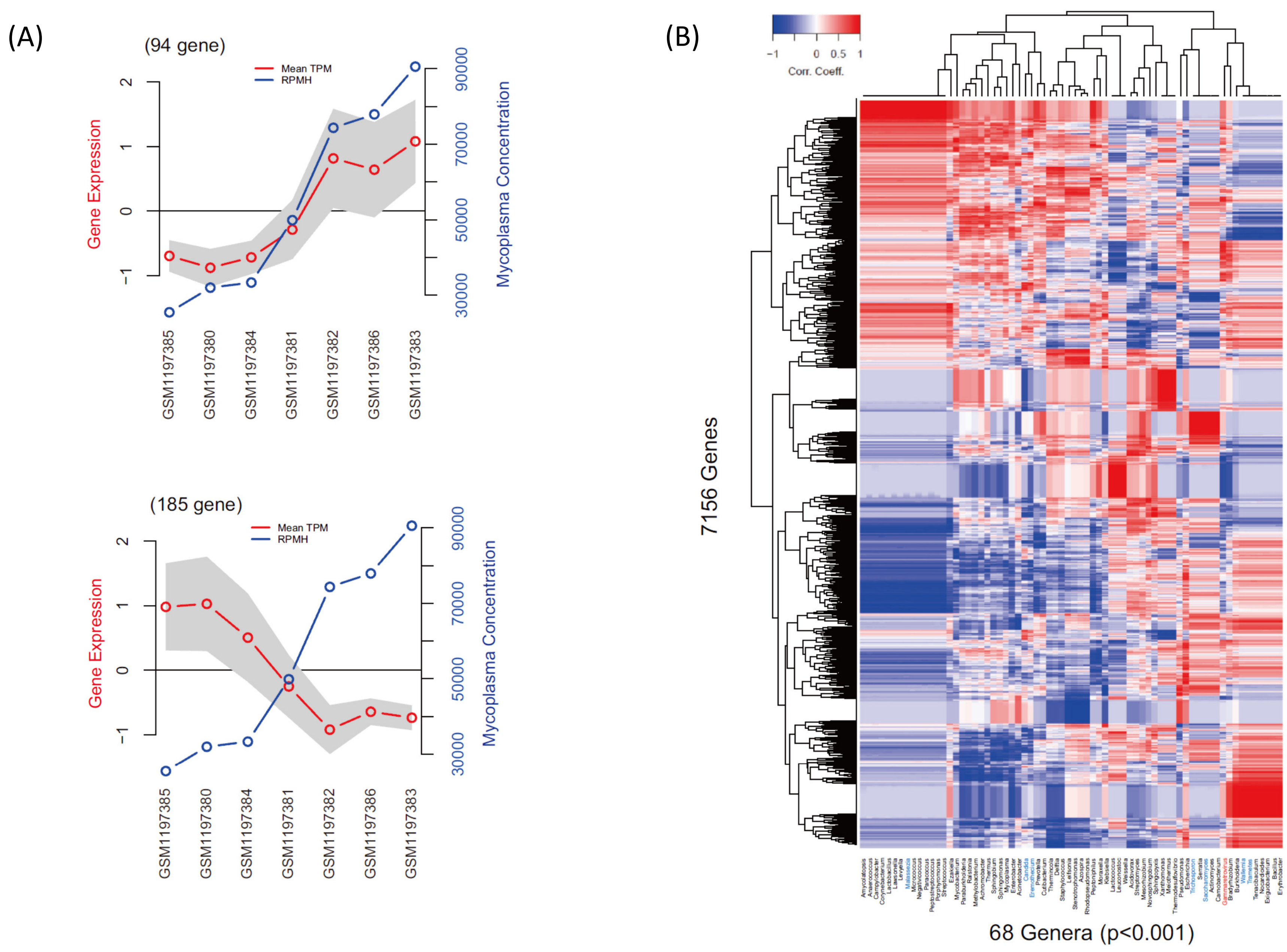

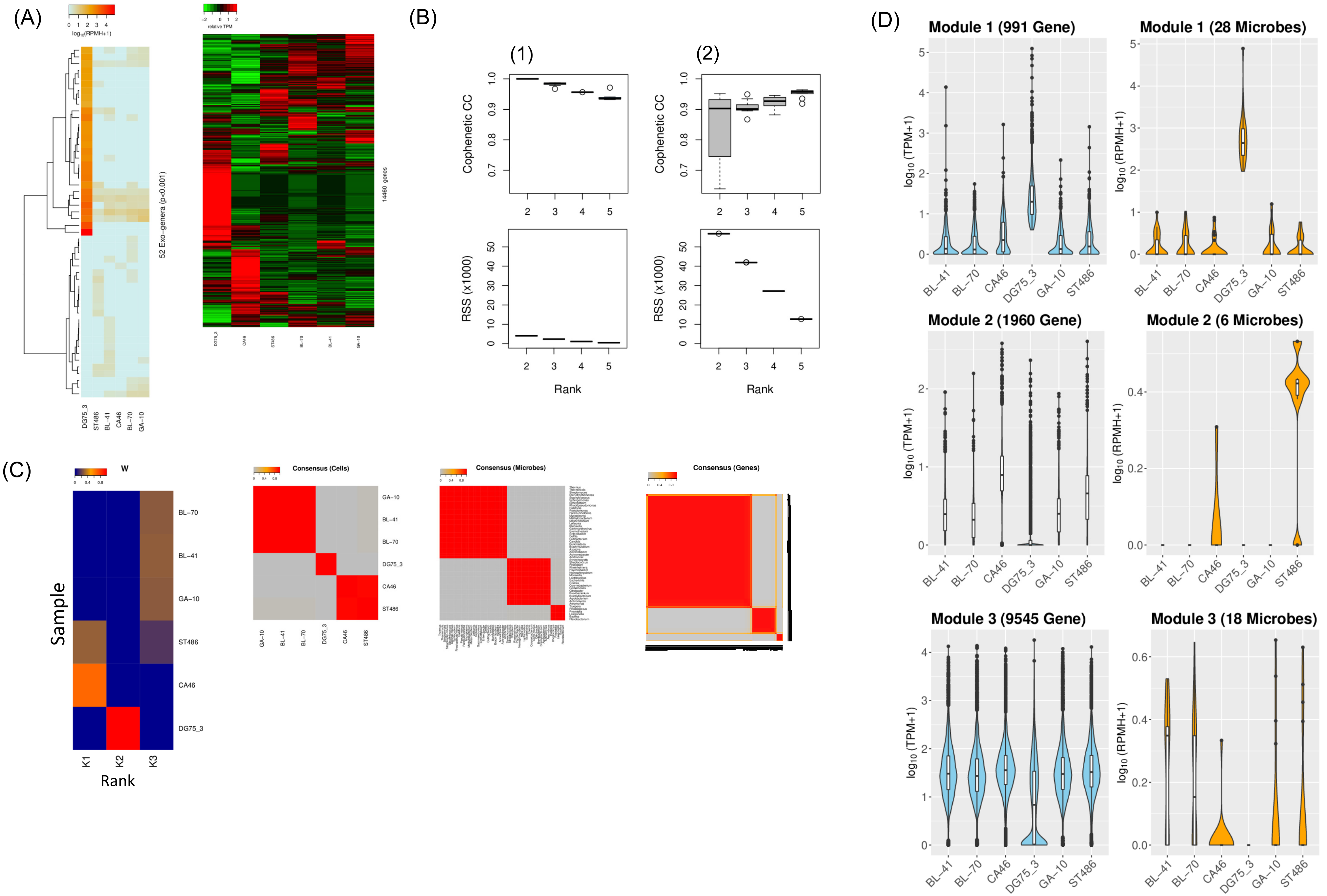
