## Additional file 2 for "A systematic NGS-based approach for contaminant detection and functional inference"

### Concept of the reversion test and its procedure for the existing pipelines

#### Concept of the reversion test

To assess the performance of NGS-based microbial contaminant detection methods, we conducted the reversion test by measuring the ratio of microbial reads that correctly map to their origin genomes. Rather than running an NGS read simulator, this test uses intact microbial 100-bp DNA fragments randomly selected from distinct species that belong to difference genera; i.e. 10,000 fragments (reads) consisting of randomly sampled 1,000 reads from each of 10 distinct species belonging to difference genera. Therefore, if the pipeline works perfectly, the species will be detected with 1,000 uniquely mapped reads from the 10 species mixture. However, this is impossible due to the higher sequence similarity among microbial species and other complex reasons.

#### Example of the reversion test

For example, 7 species of *Shigella* were selected 12 times and FDRs showed that a number of their reads were mapped to other species (i.e. TN) including *Shigella* species as shown below. These results were used in Fig. 1B (Species-level).

| FDR | TP | TN | Total | Type | Name |
|---|---|---|---|---|---|
| 0.6330 | 367 | 633 | 1000 | Species | "Shigella boydii CDC 3083-94" |
| 0.4780 | 522 | 478 | 1000 | Species | "Shigella dysenteriae Sd197" |
| 0.7830 | 651 | 2349 | 3000 | Species | "Shigella flexneri 2002017" |
| 0.5690 | 862 | 1138 | 2000 | Species | "Shigella flexneri 2003036" |
| 0.5390 | 922 | 1078 | 2000 | Species | "Shigella flexneri 5 str. 8401" |
| 0.6890 | 311 | 689 | 1000 | Species | "Shigella sonnei 53G" |
| 0.7160 | 568 | 1432 | 2000 | Species | "Shigella sonnei Ss046" |

When the TN reads mapped to *Shigella* species were counted as TP (i.e. grouping those at *Shigella* genus, level), only 1,990 out of 12,000 reads were mapped to other species not belonging to *Shigella* genus, which suggests relatively higher intra-species sequence similarity among *Shigella* species (i.e. reduced FDR to 0.17). These results were plotted in Fig. 1B (Genus-level).

| FDR | TP | TN | Total | Type | Name |
|---|---|---|---|---|---|
| 0.1658 | 10010 | 1990 | 12000 | Genus | "Shigella" |

Conceptually, in a single run of the reversion test, a species of *Shigella* appears once in the 10 species mixture (*Shigella* genus is also once) and has 1,000 reads. However, we observed that the quantifications of these species (i.e. the total mapped reads) are more than 1,000 as shown below. The quantifications consisted of uniquely mapped reads, which tend to be less than 1,000 (loss of accuracy), and reads mapped to multiple species in the input 10 species mixture (multi-mapped reads). Since the multi-mapped reads give ambiguity whether they are *Shigella* origin or not (gain of ambiguity), we penalized these multi-mapped reads using the proposed scoring scheme and included them for quantifying the species. The weighted scores show the dynamical contribution of the multi-mapped reads and their importance; that is, the inadequacy in the consideration of uniquely mapped reads only and the need for careful handling of multi-genera-hits (e.g. the case of “Sim_660”). These results were plotted in Fig. 1C.

| Total | Multi | Uniq | Weighted | Genus | Simulation |
|---|---|---|---|---|---|
| 1002 | 66 | 936 | 956.924 | Shigella | Sim_836 |
| 1003 | 101 | 902 | 939.293 | Shigella | Sim_399 |
| 1003 | 57 | 946 | 961.560 | Shigella | Sim_631 |
| 1004 | 40 | 964 | 975.921 | Shigella | Sim_66 |
| 1004 | 51 | 953 | 966.854 | Shigella | Sim_310 |
| 1007 | 86 | 921 | 942.379 | Shigella | Sim_545 |
| 1013 | 50 | 963 | 975.927 | Shigella | Sim_529 |
| 1016 | 91 | 925 | 953.424 | Shigella | Sim_103 |
| 1037 | 101 | 936 | 965.761 | Shigella | Sim_122 |
| 1063 | 293 | 770 | 870.860 | Shigella | Sim_334 |
| 1075 | 320 | 755 | 854.681 | Shigella | Sim_725 |
| 1888 | 1849 | 39 | 822.330 | Shigella | Sim_660 |

**PathSeq pipeline (BWA-MEM aligner)**

We installed the GATK4 package (ver. 4.0.5.1) and downloaded PathSeq databases (https://software.broadinstitute.org/gatk/, Accessed 12 Jun 2018). To reduce the influence of its filtering steps, a dummy host k-mer library was built with HG38 chrM and relevant parameters were turned off. The command line is given as follows;

```
%>gatk PathSeqPipelineSpark --java-options "-Xmx40G" \
    --input input_reads \
    --filter-bwa-image chrM.fa.img \
    --kmer-file chrM.hss \
    --min-clipped-read-length 20 \
    --microbe-fasta pathseq_microbe.fa \
    --microbe-bwa-image pathseq_microbe.fa.img \
    --taxonomy-file pathseq_taxonomy.db \
    --output output.pathseq.bam \
    --scores-output output.pathseq.txt \
    --skip-quality-filters true \
    --disable-tool-default-read-filters true
```

Numbers from the output file *output.pathseq.txt* were gathered; if the column ‘type’ is ‘genus’ and its ‘name’ exists in the answers, the read number in the column ‘unambiguous’ was counted as a true positive and that after subtracting the ‘unambiguous’ reads from the column ‘reads’ as a true negative. To get the distribution of mapped-read types (i.e. unique or multiple hits), the read count in the column ‘unambiguous’ was gathered as the unique-genus-hits, and the true positive reads were assumed to be multi-genera-hits.

**DecontaMiner (BLASTn aligner)**

We installed the DecontaMiner (ver. 1.4) and its databases (http://www-labgtp.na.icar.cnr.it/decontaminer/, Accessed 18 Jul 2019). The command line is given as follows;

```
%>decontaMiner.sh -s S -Q n -R n –bfv -i input_read_directory -o output_directory -c configure_file
%>filterBlastInfo.sh -i output_directory/RESULTS/BACTERIA/ -sS -m0 -g0
%>filterBlastInfo.sh -i output_directory/RESULTS/FUNGI/ -sS -m0 -g0
%>filterBlastInfo.sh -i output_directory/RESULTS/VIRUSES/ -sS -m0 -g0 –VV
%>collectInfo.sh -i output_directory/RESULTS/BACTERIA/COLLECTED_INFO/VALID/ -t 1
%>collectInfo.sh -i output_directory/RESULTS/FUNGI/COLLECTED_INFO/VALID/ -t 1
%>collectInfo.sh -i output_directory/RESULTS/VIRUSES/COLLECTED_INFO/VALID/ -t 1 -VV
```

The genus names and their 'valid' read counts reported at each run from the file '_unmapped_subject_summary_ge_CT_1.txt' in the directory 'COLLECTED_INFO/VALID/' of bacteria, fungi, and viruses were collected. If the genus exists in the answers, the genus was counted as "correct detection"; otherwise, it was counted as "false detection". To gather the uniq-genus-hits and multi-genera-hits requires careful handling because of the multiple local alignments by BLASTn. In addition, a few of the input 10,000 reads were filtered by the pipeline, resulting in different total inputs among runs. For these reasons, to get the distribution of mapped-read types, the percentages of the total inputs were calculated, rather than counting at the genus level, by using the 'valid' and 'ambiguous' read counts described at the file '_unmapped_stats.txt' in the directory 'COLLECTED_INFO/' of bacteria, fungi, and viruses.

**FastQ Screen (Bowtie2 aligner)**

The FastQ Screen (ver. 0.14.0) was installed and configured with Bowtie2 ('-k 2 --very-fast-local') (https://www.bioinformatics.babraham.ac.uk/projects/fastq_screen/, Accessed 18 Jul 2019). The command line is given as follows;

%>fastq_screen –conf *configuration_file* --outdir *output_directory* --aligner bowtie2 --subset 10000

This pipeline accepts 32 genomes as the maximum number of reference genomes. Hence, each run was configured by aligning only the 10 answer microbe genomes, which means that the statistics are based on more relaxed criteria than those of other pipelines. For example, if a input read set was prepared from the species Clostridium, Nitritalea, Dehalococcoides, Campylobacter, Sinomonas, Pseudoalteromonas, Streptomyces, Bacillus, Desulfovibrio, and Brucella, the configuration file is given by;

```
BOWTIE2 path_to_bowtie2
THREADS 4
DATABASE	Clostridium	path_to_bowtie2_index
DATABASE	Nitritalea	path_to_bowtie2_index
DATABASE	Dehalococcoides	path_to_bowtie2_index
DATABASE	Campylobacter	path_to_bowtie2_index
DATABASE	Sinomonas	path_to_bowtie2_index
DATABASE	Pseudoalteromonas	path_to_bowtie2_index
DATABASE	Streptomyces	path_to_bowtie2_index
DATABASE	Bacillus	path_to_bowtie2_index
DATABASE	Desulfovibrio	path_to_bowtie2_index
DATABASE	Brucella	path_to_bowtie2_index
```

From the output file of each run, the total number of uniq-genus-hits was collected by summing the 5th column 'One_hit_one_genome' and the 7th column 'Multiple_hits_one_genome'. Also, the total number of multi-genera-hits from the 11th column 'Multiple_hits_multiple_genomes' was collected. Finally, the uniq-genus-hits for a genus were counted as a true positive, and the read counts after subtracting the uniq-genus-hits from the total mapped reads for the genus were counted as a true negative. For example, if an output file reported as follows;

```
#Clostridium	10000	8976	89.76	0	0.00	988	9.88	5	0.05	31	0.31
```

This means that 8,976 of 10,000 input reads mapped to neither of the genomes listed in the configuration file, and 988 reads were uniq-genus-hits, and 31 reads were multi-genera-hits that mapped to several genomes. The 9th column ‘One_hit_multiple_genomes’ was ignored because we did not know which genomes were included in “multiple_genomes”.
